# An E3 ubiquitin ligase from *Nicotiana benthamiana* targets the replicase of *Bamboo mosaic virus* and restricts its replication

**DOI:** 10.1101/434076

**Authors:** I-Hsuan Chen, Jui-En Chang, Chen-Yu Wu, Ying-Ping Huang, Yau-Huei Hsu, Ching-Hsiu Tsai

## Abstract

One upregulated host gene identified previously was found involved in the infection process of *Bamboo mosaic virus* (BaMV). The full-length cDNA of this gene was cloned by 5′- and 3′-rapid amplification of cDNA ends and found to encode a polypeptide containing a conserved RING-domain and a transmembrane domain. The gene might function as an E3 ubiquitin ligase. We designated this protein in *Nicotiana benthamiana* as ubiquitin E3 ligase containing RING-domain 1 (NbUbE3R1). Further characterization by using *Tobacco rattle virus*-based virus-induced gene silencing revealed an increased BaMV accumulation in both knockdown plants and protoplasts. To further inspect the functional role of NbUbE3R1 in BaMV accumulation, NbUbE3R1 was expressed in *N. benthamiana* plants. The wild-type NbUbE3R1-orange fluorescent protein (NbUbE3R1-OFP), NbUbE3R1/△TM-OFP (removal of the transmembrane domain) and NbUbE3R1/mRING-OFP (mutation at the RING domain, the E2 interaction site) were transiently expressed in plants. NbUbE3R1 and its derivatives all functioned in restricting BaMV accumulation. The common feature of these constructs was the intact substrate-interacting domain. Yeast two-hybrid and co-immunoprecipitation experiments used to determine the possible viral-encoded substrate of NbUbE3R1 revealed the replicase of BaMV as the possible substrate. In conclusion, we identified an upregulated gene, NbUbE3R1, that plays a role in BaMV replication.

## Introduction

Plants have evolved various strategies to face rapid environmental changes and pathogen invasions (Boller and Felix, 2009; Cohn *et al.*, 2001). Recognizing and confronting these pathogens are involved in regulating the massive gene expression involved in plant-pathogen interactions (Hou *et al.*, 2009). Ubiquitin-mediated protein degradation is widely used among eukaryotes. Many ubiquitin-related genes in *Arabidopsis thaliana* have been characterized. Biochemical and proteomic studies revealed that the ubiquitin proteasome system (UPS) plays central roles in many processes of plants (Alcaide-Loridan and Jupin, 2012; Bachmair *et al.*, 2001; Vierstra, 2009). Plants exploit the UPS as a major regulatory process in hormone signaling, regulation of chromatin structure and transcription, tailoring morphogenesis, response to environmental challenges, and self-recognition and attacking pathogens (Vierstra, 2009).

In the UPS process, ubiquitin is first activated by the ATP-dependent ubiquitin-activating enzyme (E1). The activated ubiquitin is then transthiolated to ubiquitin-conjugation enzyme (E2). Ubiquitin E3 ligase (E3), the key component for UPS targeting specificity, interacts with E2-ubiquitin and the target protein to facilitate the transfer of ubiquitin and thus generate the specificity of ubiquitin modification (Metzger *et al.*, 2014; Yuan *et al.*, 2013).

On the basis of their composition and the activation mechanisms, four main types of E3 ligases are known in plants: homologous to the E6AP carboxyl terminus (HECT), really interesting new gene (RING), U-box, and cullin (a molecular scaffold protein)-RING ligases (CRLs) (Vierstra, 2009). The HECT type of E3 ligase is activated by transthiolation with ubiquitin, which is then conjugated to the substrate. RING or U-box E3 ligase mediates the transfer of ubiquitin from E2 directly to the substrate. The RING domain of E3 chelates two Zn^2+^ ions in a cross-brace structure with octet Cys and His residues used as a platform for E2 binding. RINGs are composed of one or two His residues, C3H2C3 (RING-H2) or C3HC4 (RING-HC), and other variations (Metzger *et al.*, 2014). The U-box exploits the electrostatic interaction to stabilize the ubi-E2 binding pocket (Vierstra, 2009). The CRLs consist of a cullin and a RING-containing domain, RING-BOX 1, that could bind the ubi-E2 and various adaptors and recognize target proteins (Hua and Vierstra, 2011). The target proteins have different fates depending on the type and extent of the ubiquitination.

Unusual expression levels of ubiquitin and E1 and/or E2 enzyme in the UPS pathway were demonstrated to have a comprehensive effect on cell reprogramming in the plant defense system. The E3 ligases involved in plant–pathogen and gene–gene interactions result in induction of disease resistance (Delaure *et al.*, 2008; Zeng *et al.*, 2006). Furthermore, the UPS was shown to have a resistance role in viral invasion (Takizawa *et al.*, 2005) or to be involved in specific gene-mediated resistance responses (Dielen *et al.*, 2011). For example, interfering in the ubiquitin conjugation pathway could induce different plant responses to infection with *Tobacco mosaic virus*, which supports that the UPS participates in the virus–host interaction (Alcaide-Loridan and Jupin, 2012; Becker *et al.*, 1993). The movement proteins (MPs) of *Tobamovirus*, which facilitate viral cell-to-cell transport, were found to be polyubiquitinated and degraded by 26S proteasome (Reichel and Beachy, 2000). The 66-kDa RNA-dependent RNA polymerase (RdRp) of *Turnip yellow mosaic virus* (TYMV) was targeted by the UPS in infected plant cells and downregulated viral replication (Camborde *et al.*, 2010). These findings reveal the link between the plant UPS and viral pathogens; however, the detailed mechanism is still unknown (Alcaide-Loridan and Jupin, 2012).

*Bamboo mosaic virus* (BaMV) infects more than 90% of bamboo plants. The virus induces yellow mosaic patterns on leaves and brown streaks on shoots and young culms, which causes a strong taste of shoots and results in decreased quality and output of bamboo. BaMV is believed to be the largest obstacle in the production of bamboo in Taiwan (Hsu *et al.*, 2000). BaMV, a flexuous-rod virus composed of coat protein (CP) and genomic RNA, belongs to the *Potexvirus* genus of the *Flexiviridae* family (Lin *et al.*, 1977). The genome of BaMV is 6.4 kb long and has a 5′ m^7^GpppG structure and a 3′ poly(A) tail containing five conserved open reading frames (ORFs) (Lin *et al.*, 1994). ORF1 encodes a 155-kDa polypeptide containing three functional domains: the capping enzyme domain, with GTP methyltransferase and S-adenosylmethionine(AdoMet)-dependent guanylyltransferse enzyme activities (Huang *et al.*, 2004; Li *et al.*, 2001a; Li *et al.*, 2001b); the helicase-like domain, with nucleoside triphophatase and RNA 5′-triphosphatase activities (Li *et al.*, 2001b); and a RdRp domain that recognizes the 3′-untranslated region of the BaMV genome and is involved in viral RNA replication (Huang *et al.*, 2001; Li *et al.*, 1998). ORFs 2-4 encode three overlapping MPs (termed triple gene block, TGB) for viral cell-to-cell movement (Chen *et al.*, 2012). ORF5 encodes a 25-kDa viral capsid protein required for viral RNA encapsidation (Chen *et al.*, 2012; Chen *et al.*, 2010). Two major subgenomic RNAs of approximately 2 and 1 kb were found in infected cells (Tsai *et al.*, 1999).

The defense mechanism in *Nicotiana benthamiana* was suggested to be compromised (Christie and Crawford, 1978) and susceptible to numerous viral pathogens (Bombarely *et al.*, 2012). Therefore, *N. benthamiana* plants have been exploited to mature the virus-induced gene silencing (VIGS) technology and explore RNA interference mechanisms (Angell and Baulcombe, 1997; Baulcombe, 2004; Baulcombe, 1999; Kumagai *et al.*, 1995). *N. benthamiana* is widely used as a model plant for research on plant–microbe interactions (Goodin *et al.*, 2008). Previously, our lab used the cDNA-amplified fragment length polymorphism (AFLP) technique to screen and identify BaMV infection-associated genes in *N. benthamiana* (Cheng *et al.*, 2010). VIGS was then used to knock down gene expression in *N. benthamiana* for further investigation. Among gene fragments upregulated in *N. benthamiana*, a serine/threonine kinase-like (*NbSTKL*) of *N. benthamiana* was shown to facilitate cell-to-cell movement (Cheng *et al.*, 2013). A new tau group of the GTS gene, *NbGSTU4,* was demonstrated to bind the 3′UTR of BaMV RNA and enhance viral RNA replication *in vitro* (Chen *et al.*, 2013). Among other upregulated gene fragments, *ACGT2-1* was further studied in this work.

The aim of this study was to uncover the role of *ACGT2-1* in the infection cycle of BaMV. We isolated the full-length *ACGT2-1* cDNA clone, which encodes a RING-containing protein of *N. benthamiana* and may play a key role in the ubiquitination pathway.

## Materials and Methods

### Plants and growth conditions

*N. benthamiana* plants were grown in a growth room with 16-hr day length at 28°C.

### mRNA purification and quantification

Total RNA was extracted from mock- or 500 ng BaMV virion-inoculated *N. benthamiana* plants by using Tripure isolation reagent (Roche, Germany) according to the manufacturer’s instructions. RNA of approximately 75 μg in a 100 μl-reaction was incubated at 65°C for 2 min and mixed with Dynabeads oligo(dT)_25_ for 5 min. The reaction was placed in the Dynabeads Magnetic Particle Concentrator (MPC) for 30 sec, and the supernatant was removed. The beads with poly(A)^+^ RNA were washed twice with washing buffer (10 mM Tris-HCl, pH 7.5, 1 M LiCl and 2 mM EDTA), incubated with 10 μl ddH_2_O at 80°C for 2 min, and placed in the MPC. The poly(A)-tailed mRNAs in water were transferred to a new tube.

Approximately 1 μg mRNA was mixed with 5 μM Oligo(dT)_20_ primer, incubated at 70°C for 5 min and kept on ice. The reaction contained the pre-incubated mRNA, 3 mM MgCl_2_, 0.5 mM dNTP, 6 Us RNaseOUT (RNase inhibitor, Invitrogen, Carlsbad, CA, USA), and 5 Us reverse transcriptase (Promega, Madison, WI, USA).

To confirm the expression profile of ACGT2-1 after BaMV infection, primer sets GT2-1/F1 (5′TAAGTGTAAACATGATGTTCCAAAT3′) and GT2-1/R1 (5′CGAACCTATA ACCTCCTGAT3′) and actin F (5′GTGGTTTCATGAATGCCAGCA3′) and actin R (5′GATGAAGATACTCAC AGAAAGA3′) were used in PCR to detect the expression of *NbUbE3R1* and *β-Actin*, respectively.

The expression of *NbUbE3R1* was analyzed by qRT-PCR with the KAPA SYBR FAST qPCR Kit Master Mix (KapaBiosystems, Boston, MA, USA). The expression of actin was used for normalization. The reactions were set at thermal cycling conditions with 95°C for 3 min, and 40 cycles at 95°C for 3 sec followed by 60°C for 20 sec. The specific primers for quantitative RT-PCR were GT2-1/F1 and GT2-1/R4 (5′GTTCTTGAG GAATACAAAGTGCAC3′).

### Constructs

The cDNA fragment (*ACGT2-1*) (Cheng *et al.*, 2010) was released from pGEM-T Easy vector (Promega) by digestion with *Eco*RI and cloned into the TRV2 vector for knockdown experiments. 3′ rapid amplification of cDNA ends (RACE) involved reverse transcription of mRNAs isolated from healthy *N. benthamiana* plants by using the primer 3′RACE/RT (5′CAACTCGAGCCCGGGATCCCTAT_25_NN3′). Amplification involved two PCR reactions with one forward primer GT2-1/F1 and two reverse primers: 3′RACE/R1 (5′CAACTCGAGCCCGGGAT 3′) and 3′RACE/R2 (5′GAGCCCGGGATCCCTAT3′). For 5′ RACE, the first-strand cDNAs were synthesized by using the SMARTer RACE cDNA Amplification Kit (Clontech Laboratories, Mountain View, CA, USA) according to the manufacturer’s protocol. Primers 5′RACE/long (5′CTAATACGACTCACTATAGGGCAAGCAGTGGTATCAACGCAGAGT3′) and 5′RACE/R1 (5′CAGCCCAGTGCACTAAGCTCCCGCTATATGCGTGGTCCG3′) were used in PCR. The products were then diluted 10-fold and used as the template. The primer set 5′RACE/short (5′CTAATACGACTCACTATAGGGC3′) and 5′RACE/R2 (5′GGGCCAGATCACAAGGGTTTATTGTACG3′) was used in the second round of PCR. The final products were gel-purified and cloned into the pGEM-T Easy vector (Promega).

The ORF of *NbUbE3R1* was PCR-amplified with the primer set GT2-1/F5 (5′GG<u>TCTAGA</u>ATGGCCATTTTTGATAG3′) (*Xba*I site underlined) and GT2-1/R5 (5′G<u>GTACC</u>CATTGTTTGTCTTAGATCTT3′) (*Kpn*I site underlined). The amplified DNA fragment was cloned into the pEpyon/mOrange2 vector with *Xba*I and *Kpn*I sites to create a construct that produced the fusion protein NbUbE3R1-orange fluorescent protein (NbUbE3R1-OFP). The mutant construct with removal of the transmembrane domain NbUbE3R1/∆TM-OFP was created by PCR with the primer set GT2-1/F4 (5′G<u>TCTAGA</u>ATGGTTGATCATCCAATTTGGT3′) (*Xba*I site underlined) and GT2-1/R5. The amplified fragment was cloned into the pGEM-T Easy vector and transferred to the pEpyon/mOrange2 vector with *Xba*I and *Kpn*I sites after sequence verification.

The mutant NbUbE3R1/mRING-OFP was created by three cloning steps. First, the ORF of NbUbE3R1 was amplified with two sets of primer pairs, ME3F1 (5′GCGCTAAGATCACACAAGAATTGCGCAGCGTGTCGCGCTCC3′)/ME3R1 (5′G<u>CTCGAG</u>ATGAATTATTTTAGCAGCTGAAGAATCC3′) (*Ava*I underlined) and ME3F2 (5′GGTATA<u>TACGTA</u>CAATAGGACTATCACA3′) (*Sna*BI underlined)/ME3R2 (5′TTCAGTCAAACAAGCAGCACAATCACTTCC3′). The PCR products generated were 389 and 118 bp, respectively. Second, the 118-bp product was used as a megaprimer together with the primer ME3R3 (5′CGCTGCGCAATTCTTGTGTGATCTTAGCGCAGTATCAATACAAGG3′) to generate a 223-bp DNA fragment. This PCR product overlapped with the 389-bp PCR product by 30 bp. Finally, the two overlapped DNA fragments (389 and 233 bp) were used to synthesize a 582-bp DNA fragment by overlapped extension PCR. The final PCR product containing five point mutations (Fig. S2) was cloned into the pGEM-T easy vector and verified. The mutant DNA fragment was substituted for the original wild-type DNA fragment with *Sna*BI and *Ava*I sites. Finally, the mutant NbUbE3R1/mRING was subcloned into the pEpyon/mOrange2 vector.

For yeast two-hybrid assay, the coding sequence of NbUbE3R1/∆TM was amplified with the primer pair pYES-KpnI (5′<u>GGTACC</u>ATGGTTGATCATCCAATTTGGTATATACGTACAATAG3′) (*Kpn*I underlined) and pYES-SacI (5′<u>GAGCTC</u>CATTGTTTGTCTTAGATCTT3′) (*Sac*I underlined) and cloned into the pGEM-T Easy vector. The coding sequence was then subcloned into the *Kpn*I/*Sac*I sites of pYESTrp2 to generate the prey plasmids.

### Knockdown experiment and virus inoculation

TRV-based VIGS was used in the gene-knockdown experiment. The cDNA fragment (ACGT2-1) derived from cDNA-AFLP was cloned into the pTRV2 vector. The TRV2 vectors containing the phytoene desaturase gene and luciferase gene were used as silencing and negative controls, respectively. *Agrobacterium tumefaciens* C58C1 strain containing pTRV1 or pTRV2 and its derivatives was cultured at 30°C with the addition of 10 mM MES pH 5.6 and 6 μM acetosyringone to OD_600_=1. The bacteria were pelleted and suspended into the induction buffer (10 mM MgCl_2_, 10 mM MES, 150 μM acetosyringone). The bacteria containing TRV1 and TRV2 were mixed in a 1:1 ratio for infiltration. In the knockdown experiment, 4-week-old plants were used for *Agrobacterium* infiltration with a syringe without a needle. The prepared *Agrobacterium* mixtures were infiltrated onto three leaves above the cotyledons of each plant. At 12 days post-infiltration, the fourth leaf above the infiltrated leaves was inoculated with 500 ng BaMV viral particle.

### Western blot assay

Total proteins were extracted with extraction buffer (5.6 mM Tris-HCl pH 6.8, 10.5% glycerol and 2.1% SDS), separated on 12% SDS-PAGE and transferred to nitrocellulose membrane, which was probed with the primary antibody (rabbit anti-BaMV CP), then fluorescent-labeled secondary antibody (anti-rabbit IgG, ROCKLAND). Finally, the fluorescent signal on the membrane was visualized and quantified by using LI-COR Odyssey (LI-COR Biosciences). The CP accumulation was normalized to that of Rubisco large subunit (rbcL) stained with Coomassie blue.

### Protoplast isolation and viral RNA inoculation

The silenced leaves at 14 days post agroinfiltration collected for analysis of specific gene knockdown were sliced into thin strips. Leaves were digested with enzyme solution (0.1% bovine serum albumin, 0.6 mg/ml pectinase and 12 mg/ml cellulose in 0.55 M mannitol-MES pH 5.7) in the dark at 25°C for 10-12 hr. The extracted protoplasts were filtered by using a miracloth and separated by centrifugation at 300 rpm for 7 min (KUBOTA KS-5000). Healthy protoplasts were further isolated as described (Chen *et al.*, 2018). Approximately 2.5×10^5^ protoplasts were inoculated with 1 μg BaMV viral RNA in 20% polyethylenglycol (PEG) 6000. Inoculated protoplasts were resuspended in culture medium and incubated at 25°C for 24 hr under constant light. Total protein and RNA were extracted according to previously described protocols.

### Northern blot assay

Approximately 1 μg RNA was mixed with 10 mM phosphate buffer, pH 7.0, 50% DMSO, 1 M glyoxal in a final 12 μl-reaction and incubated at 50°C for 1 hr. After denaturation, RNA was size-fractionated in an 1% agarose gel and blotted onto a nylon membrane (Hybond-N+, GE Healthcare, UK) by using 0.2 M NaOH transferring solution for 40 min. RNA was cross-linked to the membrane by UV exposure (1200J, Stratagene, USA). The membrane was prehybridized in 10 ml hybridization buffer (1x SET, 1% sodium pyrophosphate, 0.6% SDS, 10x Denhard’t, salmon sperm DNA 75 μl) at 65°C for 2 hr, then hybridized with [α-^32^P]UTP-labeled probe (10^6^ cpm) annealed to the 3′-end approximately 600 nt of BaMV RNA at 65°C overnight. The membrane was washed with washing buffer (0.5x SET, 0.1% SDS, 0.1% sodium pyrophosphate) three times for 20 min each at 65°C.

### Transient expression in N. benthamiana

*Agrobacterium* containing NbUbE3R1-OFP, NbUbE3R1/∆TM-OFP, NbUbE3R1/mRING-OFP, OFP or the silencing suppressor HcPro was cultured to OD_600_ =1. Five-week-old *N. benthamiana* plants were infiltrated with *Agrobaterium* containing the desired plasmid and that with HcPro in a 1:1 ratio. At 4 hr post-agroinfiltration, the infiltrated leaf was inoculated with 200 ng BaMV viral particles. At 3 dpi, protein was extracted from inoculated leaves for western blot analysis.

### Subcellular localization with confocal microscopy

Fluorescence images of *N. benthamiana* leaf and protoplasts expressing OFP and OFP-fusion proteins were examined by confocal laser scanning microscopy (Olympus Fluoview FV1000, Olympus, Japan) obtained at 543 nm excitation and analyzed and/or merged by using Olympus FV10-ASW1.3 viewer software.

### Bioinformatics analysis

To inspect gene-specific primers, the sequence from cDNA-AFLP was searched in the *N. benthamiana* sol genome sequence database (http://sydney.edu.au/science/molecular_bioscience/sites/benthamiana/). Web analysis was used to obtain the features of NbUbE3R1. SMART (http://smart.embl-heidelberg.de/) was used to evaluate the content of the protein, and the Aramemnon program (http://aramemnon.botanik.uni-koeln.de/request.ep) was used to predict the subcellular localization. The sequences were aligned at the Biology Workbench website (http://seqtool.sdsc.edu/).

### MG132 treatment

To examine whether NbUbE3R1 was degraded via the 26S proteasome pathway, the 26S proteasome inhibitor MG132 dissolved in DMSO was used at 50 μM. The leaves of *N. benthamiana* were infiltrated with *Agrobacterium* containing the desired plasmid. The plants expressing OFP alone, NbUbE3R1-OFP, NbUbE3R1/∆TM-OFP or NbUbE3R1/mRING-OFP were further treated with MG132 for 12 hr before sample collection. After 3 days post infiltration, total protein was extracted from inoculated leaves and examined by western blot analysis.

### Yeast two-hybrid interactions

The yeast Hybrid Hunter system (Invitrogen) was used in the study. An interaction between bait and prey protein in the system would activate the expression of the reporter gene (*his3* and *lacZ*) in *Saccharomyces cerevisiae* strain L40. The gene fragment encoding NbUbE3R1/∆TM was constructed into the prey plasmid pYESTrp2 and pLEX, designated pYES-E3/∆TM and pLEX-E3/∆TM. The replication-related DNA fragments were constructed into the bait plasmid pHybLex/Zeo and designated pLEX-Capping, -RdRp, and -Helicase. The MPs and CP genes were constructed into the prey plasmid pYESTrp2 and designated pYES-TGBp1, -TGBp2, -TGBp3, and -CP (Huang *et al.*, 2017b). The bait plasmids (pLEX-Capping, -RdRp, or -Helicase) and prey plasmid (pYES-E3/∆TM) or prey plasmids (pYES-TGBp1, -TGBp2, -TGBp3, or -CP) and bait plasmid (pLEX- E3/∆TM) were co-transformed into *S. cerevisiae* strain L40 and selected on Trp^-^/His^-^/Zeocin agar plates.

### Co-immunoprecipitation

*Agrobacterium* containing NbUbE3R1/∆TM-OFP or OFP alone mixed with the one containing the BaMV RdRp in a 1:1 ratio was infiltrated onto *N. benthamiana* leaves in the presence of HcPro. After 3 days post infiltration, total protein was extracted from plant leaves by using the extraction buffer containing 20 mM Tris-HCl, pH 7.5, 2 mM MgCl_2_, 300 mM NaCl, 0.5% NP-40, 5 mM dithiothreitol (DTT) and EDTA-free Protease Inhibitor Cocktail (Roche). The extracts were centrifuged at 4000×g at 4°C for 10 min, and supernatant was subjected to immunoprecipitation by adding 20 μl anti-OFP magnetic beads and rotating with rotamixer for 4 hr at 4°C. The beads were then washed four times with binding buffer (20 mM Tris-HCl, pH 7.5, 2 mM MgCl_2_ and 300 mM NaCl), then washed with TBS once before 4x sample buffer was added to release proteins. Finally, samples were subjected to western blot analysis with anti-HA or anti-OFP antibody.

## Results

### The gene containing ACGT2-1 is a putative C3H2C3-type zinc finger ubiquitin E3 ligase

The expression profile of the upregulated gene containing the ACGT2-1 cDNA fragment identified by cDNA-AFLP (Cheng *et al.*, 2010) was further confirmed by RT-PCR. The expression of the gene containing ACGT2-1 was upregulated at 5 to 7 dpi after BaMV inoculation (Fig. S1).

To identify the gene containing ACGT2-1 cDNA fragment, 3′ and 5′ rapid amplification of cDNA ends (RACE) was performed. The full-length cDNA obtained from RACE was about 1.7 kb. The coding region was 1218 bp and encoded a 406-amino acid polypeptide (Fig. S2). The sequence of the *in silico*-translated polypeptide was predicted to contain a transmembrane domain, a RING zinc-finger domain and a C-terminal coiled-coil region by domain architecture analysis (Fig. S2). The results derived from the sequence comparison by using the available databases indicated that the RING finger domain is highly conserved, commonly found in E3 ubiquitin ligases, and plays a key role in the ubiquitination pathway (Guzman, 2012). Therefore, the upregulated gene containing the ACGT2-1 fragment could be a putative C3H2C3-type zinc finger E3 ubiquitin ligase. We then designated this polypeptide ubiquitin E3 RING type ligase 1 of *N. benthamiana*, *NbUbE3R1*.

### The upregulated gene NbUbE3R1 is involved in BaMV infection

To investigate whether *NbUbE3R1* is involved in the BaMV infection cycle, we used TRV-based VIGS to knock down its expression in *N. benthamiana* plants. To rule out the possibility of an off-target by using the ACGT2-1 fragment to knock down the gene expression, the sequence of ACGT2-1 was analyzed in the *N. benthamiana* sol genome sequence database (Fernandez-Pozo *et al.*, 2015). We did not find any sequence matching that of ACGT2-1 with consecutive residues longer than 20 nt besides the target gene, *NbUbE3R1*. Therefore, using the sequence of ACGT2-1 to knock down the expression of *NbUbE3R1* in *N. benthamiana* could be specific.

At approximately 10 to 14 days post agroinfiltration, the morphology of *NbUbE3R1*-knockdown plants showed no obvious difference from luciferase (Luc)-knockdown plants (Fig. S3). The knockdown efficiency measured by real-time RT-PCR was reduced 27% in *NbUbE3R1*-knockdown plants as compared with control plants (Fig. 1A). The accumulation of BaMV CP in *NbUbE3R1*-knockdown plants quantified at 5 dpi was significantly increased 1.34-fold that of control plants (Fig. 1B). Therefore, the target gene was involved in the infection cycle of BaMV and possibly plays a defense role against BaMV infection.

**Fig. 1.**
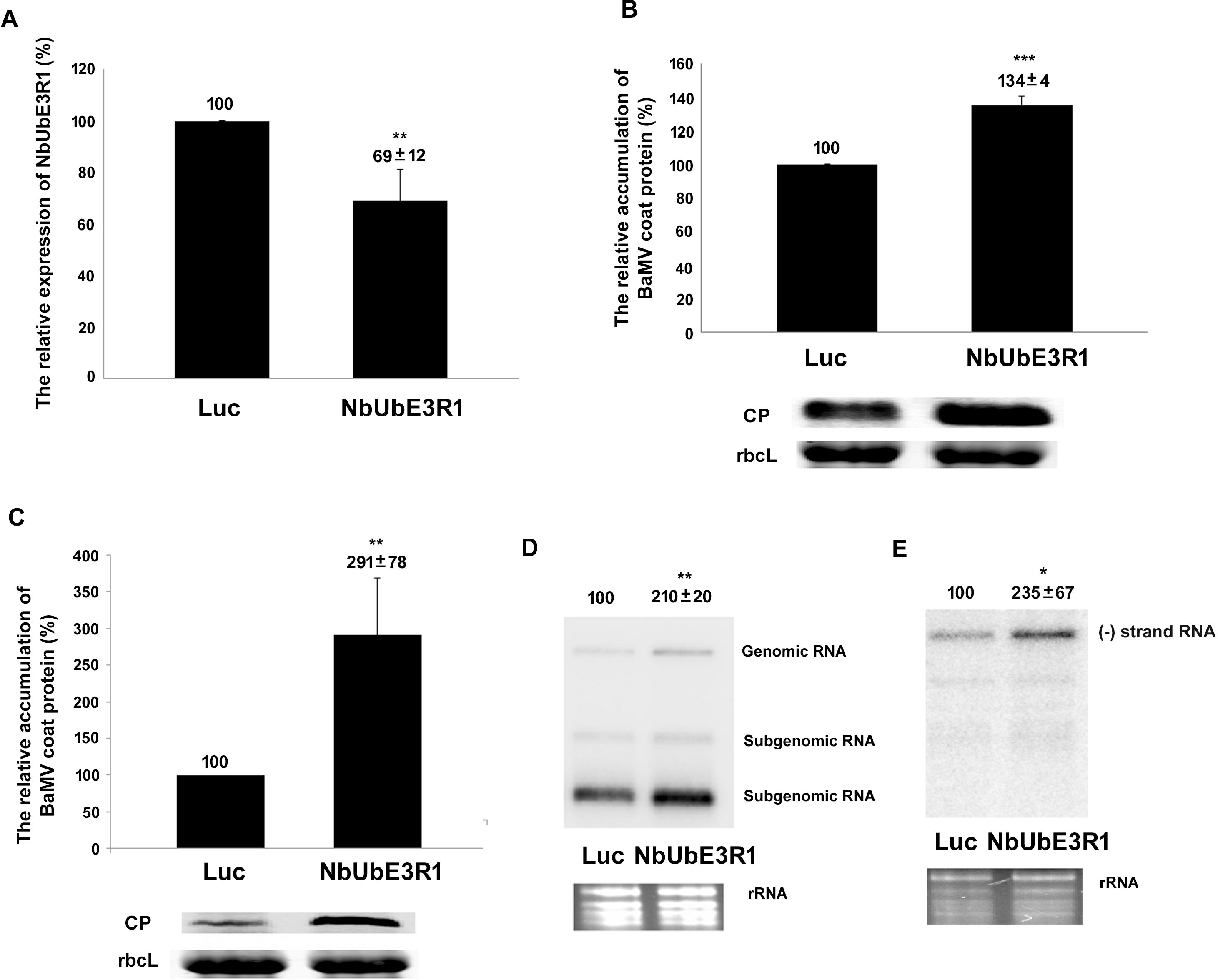
Relative expression and accumulation of *NbUbE3R1* and BaMV in *NbUbE3R1-*knockdown *Nicotiana benthamiana*. (A) Relative expression of *NbUbE3R1* in plants determined by real-time RT-PCR. The expression of actin was used for normalization. (B) The relative accumulation of BaMV coat protein in plants harvested at 5 days post inoculation (dpi) determined by western blot analysis. The expression of RuBisCo large subunit (rbcL) was the loading control. Protoplasts were isolated from *Luciferase*- or *NbUbE3R1*-knockdown plants and inoculated with BaMV viral RNA. The accumulation of BaMV coat protein (C) and viral RNA, plus-strand RNA (D) and minus-strand RNA (E), in plants harvested at 24 hr post inoculation quantified by western and northern blot analyses. The expression of RuBisCo large subunit (rbcL) and the ribosome RNA (rRNA) was the loading control in western and northern blot analyses, respectively. The expression in *Luciferase*-knockdown plants was set to 100%. The number above the statistical bar or above the blots is the mean±SE of three independent experiments. \**P*<0.05, \*\**P*<0.01, and \*\*\**P*<0.001 by Student’s *t*-test.

### NbUbE3R1 is involved in BaMV RNA replication

To inspect how NbUbE3R1 affects the accumulation of BaMV in *N. benthamiana*, protoplasts derived from the *NbUbE3R1*-knockdown plants were isolated and inoculated with viral RNA. The accumulation of BaMV CP was increased to 291% of that of control protoplasts at 24 hr post-inoculation (Fig. 1C). The accumulation of viral RNA in *NbUbE3R1*-knockdown protoplasts was significantly increased 2.10- and 2.35-fold for the plus- and minus-strand RNA, respectively, that of control protoplasts (Fig. 1D and 1E). These results suggest that the ACGT2-1-containing gene NbUbE3R1 might be involved in viral RNA synthesis.

### Subcellular localization of NbUbE3R1

Because NbUbE3R1 contains a transmembrane domain, it is likely a membrane-associated protein. Therefore, we further analyzed NbUbE3R1 in the plant membrane protein database Aramemnon to predict its possible role and found that NbUbE3R1 could be involved in the secretory pathway. To experimentally determine the subcellular location of NbUbE3R1, orange fluorescent protein (OFP)-tagged NbUbE3R1 or OFP alone was transiently expressed in *N. benthamiana* leaf and protoplasts. On confocal microscopy, wild-type NbUbE3R1-OFP clustered to form speckled spots in protoplasts (Fig. 2), whereas localization of the mutant with the N-terminus transmembrane truncation, NbUbE3R1/∆TM-OFP, was similar to that with OFP alone.

**Fig. 2.**
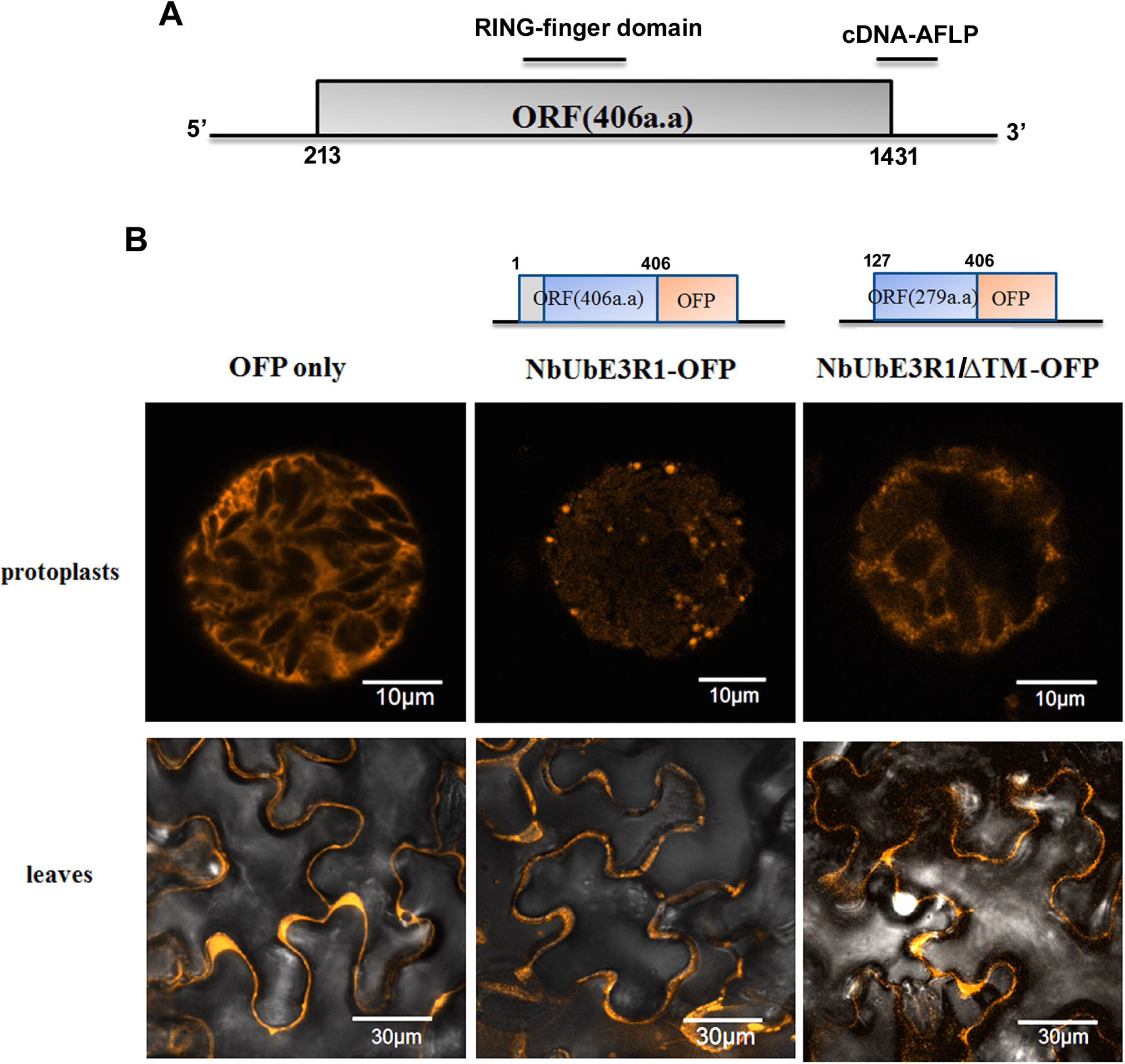
Structure diagram of the full-length cDNA of NbUbE3R1 and its subcellular localization. (A) Illustration of the cDNA of NbUbE3R1. The open reading frame (nt 213 to 1431), RING-finger domain and cDNA fragment identified from AFLP used for virus-induced gene silencing are indicated. (B) The expression of orange fluorescent protein (OFP)-fused NbUbE3R1 or NbUbE3R1/∆TM in *N. benthamiana* protoplasts (top 3 panels) or leaves (bottom 3 panels).

### NbUbE3R1 negatively regulates BaMV infection

The results from *NbUbE3R1*-knockdown experiments suggested that NbUbE3R1 could have a defensive role against BaMV. To verify this hypothesis, NbUbE3R1-OFP and its derivatives were expressed in *N. benthamiana* leaves and BaMV was inoculated in NbUbE3R1-expressed leaves. The accumulation of BaMV CP was reduced to 17% of that of control plants expressing OFP only (Fig. 3). To determine whether the membrane association of NbUbE3R1 is critical for its defensive role, we expressed NbUbE3R1/∆TM-OFP in leaves. The accumulation of BaMV CP was significantly reduced to 31% of that of the control. Therefore, NbUbE3R1 without the transmembrane domain could still have a significant role against BaMV. However, the wild-type NbUbE3R1 containing the transmembrane domain was more efficient in targeting BaMV.

**Fig. 3.**
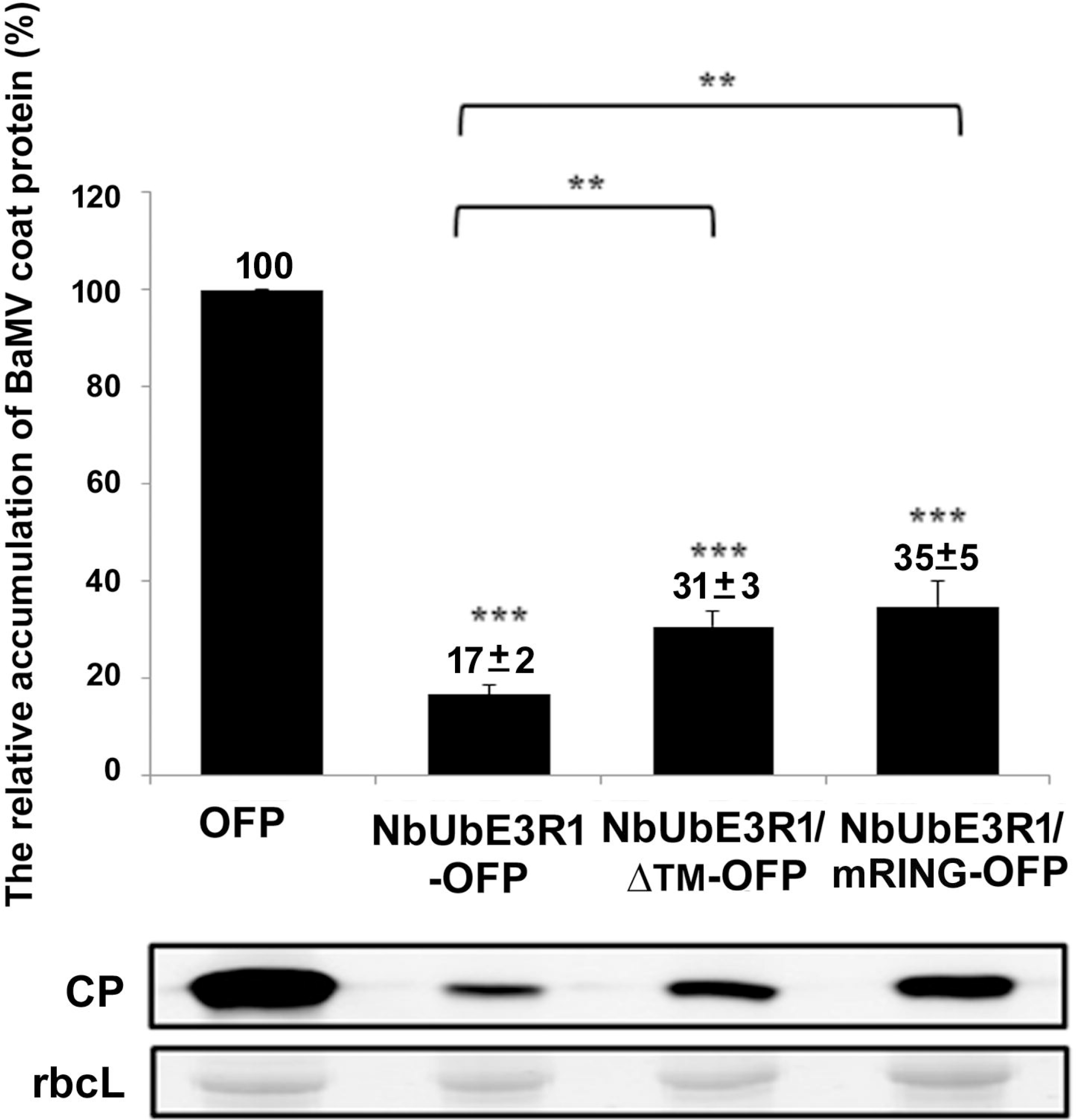
The relative accumulation of BaMV coat protein (CP) in plants expressing NbUbE3R1 or its derivatives. The accumulation of BaMV CP was quantified by western blot analysis. Total protein was extracted from 200 ng BaMV-inoculated *N. benthamiana* plants at 3 days post inoculation (dpi) transiently expressed with OFP, NbUbE3R1-OFP, NbUbE3R1/∆TM-OFP, or NbUbE3R1/mRING-OFP. The numbers above the bar are the mean±SE CP level from three independent experiments. rbcL: RuBisCo large subunit was a loading control. ** *P*<0.01, \*\*\**P*<0.001 by Student’s *t*-test.

Because NbUbE3R1is predicted to be an E3 ubiquitin ligase, the RING domain may contain an E2 binding site required for activity to target the substrate for degradation. To validate this hypothesis, a mutant with five point mutations (S166A, V167A, W196A, P204A, and L205A) in the RING finger domain of NbUbE3R1 was constructed (Fig. S2) and designated NbUbE3R1/mRING. These residues in the RING domain of E3s were reported to be potential E2-contact points (Deshaies and Joazeiro, 2009). NbUbE3R1/mRING carrying these mutations is expected to be defective in RING-E2 interaction to a certain extent, so the association of NbUbE3R1 catalytic activity and BaMV replication can be revealed. The accumulation of BaMV in NbUbE3R1/mRING-OFP-expressed leaves was approximately 35% of that of the control (Fig. 3). Therefore, NbUbE3R1 restricting BaMV replication does not need to have the full function of E3 ligase. Similarly, the reduced BaMV replication of NbUbE3R1/mRING-OFP was significantly less than that of the wild type. Thus, the transmembrane and RING finger domains of NbUbE3R1 could also play in part in negative regulation of BaMV infection.

### NbUbE3R1 protein is unstable in vivo

Transiently expressed NbUbE3R1-OFP and NbUbE3R1/mRING-OFP were barely detected by western blot analysis (Fig. 4) perhaps because of the instability of NbUbE3R1, which might undergo auto-ubiquitination or be targeted by an endogenous degradation pathway (de Bie and Ciechanover, 2011). To clarify whether the low expression of NbUbE3R1 was due to the instability from the auto-ubiquitination of NbUbE3R1, plants were treated with the 26S proteasome inhibitor MG132 after transient expression of NbUbE3R1 and its derivatives. The expression of NbUbE3R1 was elevated by MG132 treatment (Fig. 4). Also, the expression of NbUbE3R1/mRING was higher than that of NbUbE3R1 perhaps because NbUbE3R1/mRING lost its ubiquitination activity and could not undergo self-regulated auto-ubiquitination within the plant cell.

**Fig. 4.**
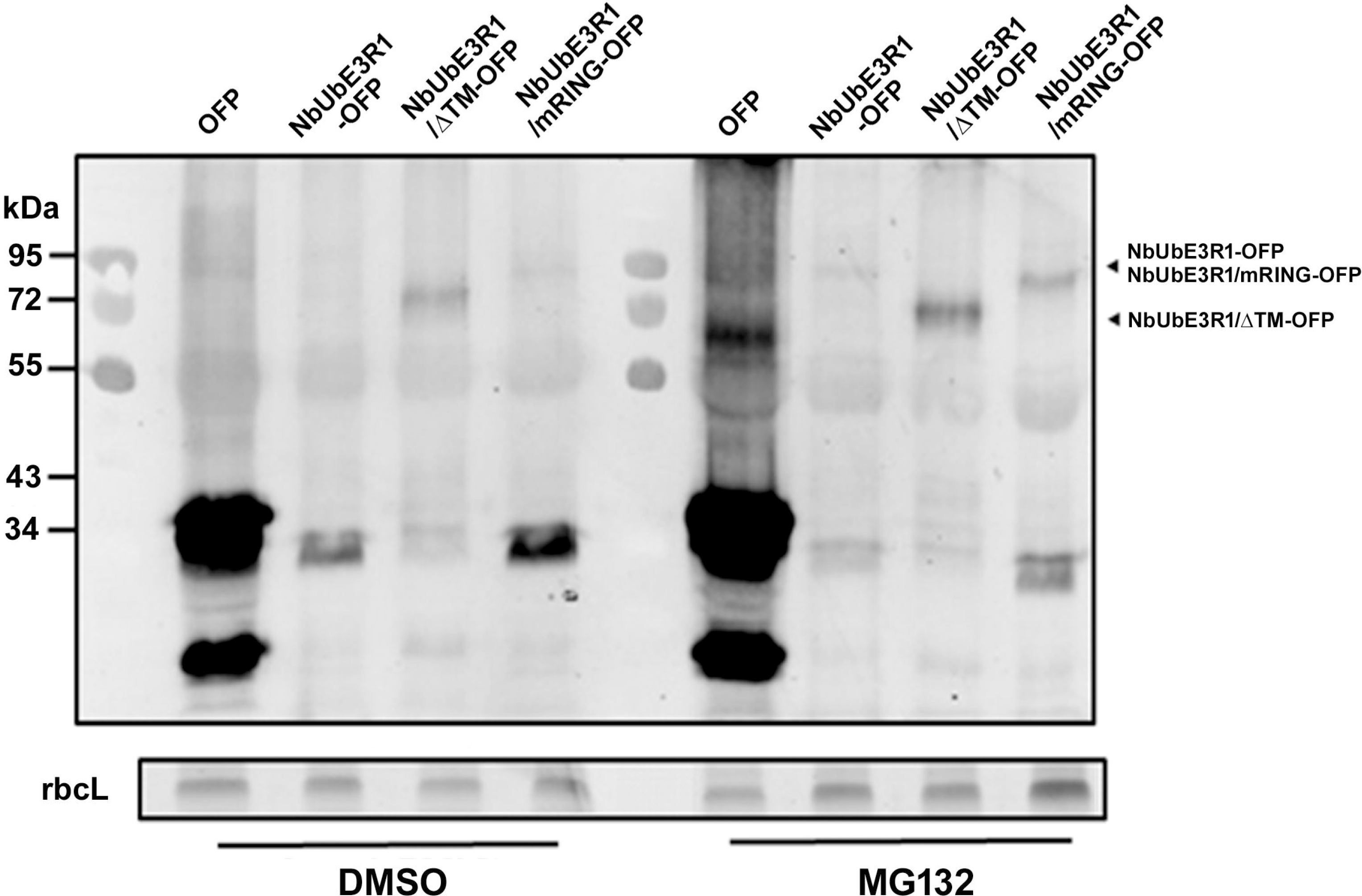
Effect of MG132 on the stability of NbUbE3R1 and its derivatives *in vivo.* The proteosome inhibitor MG132 (50 μM) or DMSO (control) was infiltrated into plants which were agroinfiltrated with OFP, NbUbE3R1-OFP, or its derivatives at 12 hr before harvesting. Total protein extracted from control and MG132-treated plants at 3 days post infiltration were subjected to western blot analysis. rbcL: Rubisco large subunit was the loading control.

### NbUbE3R1 interacts with BaMV replicase

The results from the transient expression experiments (Fig. 3) indicated that NbUbE3R1/mRING without the E3 ligase activity could still suppress BaMV replication, perhaps because the NbUbE3R1/mRING could still interact with the substrate (the BaMV replication-related protein). To elucidate the possible substrate of NbUbE3R1, we used a yeast two-hybrid assay to test whether NbUbE3R1 could interact with viral replicase. Each domain of the BaMV-encoded replicase was cloned into the pLEXA as a bait and designated pLEX/Capping, -/Helicase, and -/RdRp. The coding sequence of NbUbE3R1/∆TM (a soluble form and with suppression activity) was cloned into pYESTrp2 as a prey and then transformed into yeast containing each domain of the viral replicase. Double transformants (yeast with both pLEX and pYES) were subjected to strong selective conditions. NbUbE3R1/∆TM could interact with the RdRp domain in the yeast cells (Fig. 5).

**Fig. 5.**
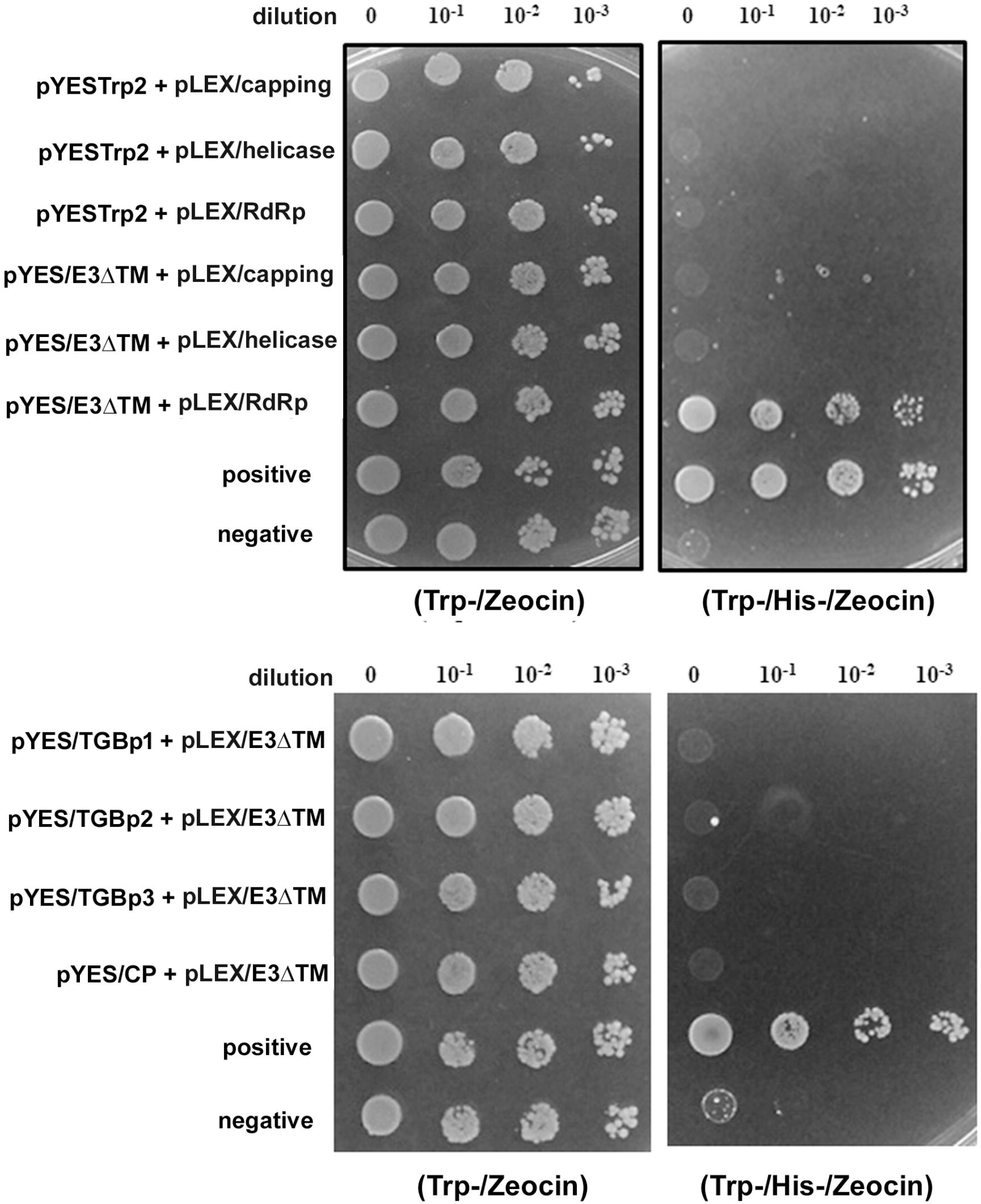
Interaction of NbUbE3R1/∆TM with BaMV replicase in yeast cells. Yeast strain L40 co-transformed with the indicated plasmids was subjected to 10-fold serial dilution and incubated with minimal medium lacking tryptophan and histidine supplemented with 3-AT (5 mM) to identify protein interactions. Yeast containing pYES-Hsp90 and pLEX-RdRp was a positive control; yeast containing the vector pHybLex/Zeo and pYESTrp2 was a negative control.

Co-immunoprecipitation was used to validate the results of the yeast two-hybrid experiments. Agroinfiltrated *N. benthamiana* leaves were used to examine the interaction between NbUbE3R1 and BaMV replicase in planta. NbUbE3R1/∆TM-OFP or OFP (a negative control) was co-infiltrated with BaMV/Rep-HA, an infectious cDNA clone with the replicase tagged with HA at its C-terminus. Total proteins were immunoprecipitated with anti-OFP magnetic beads. The replicase of BaMV could co-precipitate with NbUbE3R1/∆TM (Fig. 6).

**Fig. 6.**
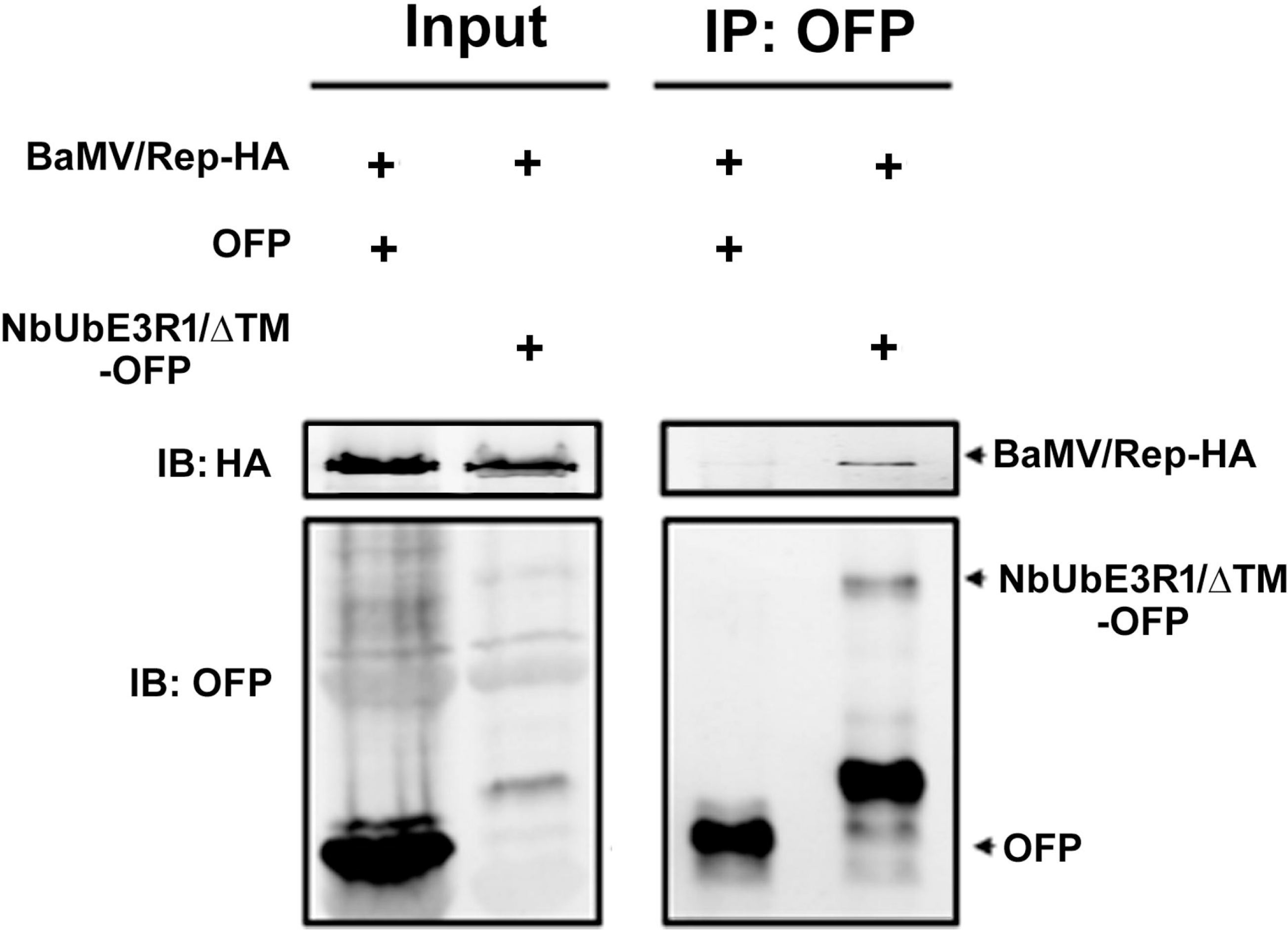
Co-immunoprecipitation analysis of the interaction between NbUbE3R1/∆TM and BaMV replication protein *in vivo*. Total protein (input) was extracted from *N. benthamiana* leaves co-infiltrated with *A. tumefaciens* cells harboring BaMV/Rep-HA and OFP only or NbUbE3R1/∆TM as indicated. An equal volume of input proteins was then incubated with anti-OFP magnetic beads. Samples before (Input) and after (IP) immunoprecipitation were subjected to western blot analyses (IB) with antibodies against HA or OFP.

## Discussion

The study of differentially expressed genes after virus infection is a strategy for identifying host factors involved in virus infection cycles. We used a cDNA-AFLP technique to identify a number of these genes (Cheng *et al.*, 2010) and revealed their involvement in BaMV infection (Huang *et al.*, 2017a). One of the host factors in this study was demonstrated to play a defensive role against BaMV replication. Sequence analysis indicated that this gene contains a C3H2C3-type RING finger domain and is possibly an E3 ubiquitin ligase. The structure of the RING-H2 domain in NbUbE3R1 is similar to that of Arabidopsis Tóxicos en Levadura 54 (ATL54), which was demonstrated to harbor *in vitro* ubiquitin ligase activity (Noda *et al.*, 2013). Therefore, we proposed that NbUbE3R1 could mediate a degradation or modification of BaMV replication-related protein.

Bioinformatics analysis indicated that NbUbE3R1 could be in the secretory pathway. We used OFP-tagged NbUbE3R1 to inspect the subcellular localization. However, the expression of this protein was so low and barely detected on western blot analysis perhaps because NbUbE3R1 was unstable and might undergo auto-ubiquitination and self-destruction (Lin *et al.*, 2008). We used MG132, a proteasome inhibitor, to treat leaves before infiltration, but we still could not obtain strong fluorescent signals. The expression experiments indicated that NbUbE3R1 associated with membrane or not could target BaMV and reduce its accumulation. However, NbUbE3R1 targeting to the membrane could be critical and more efficient against BaMV infection. Mutant NbUbE3R1/∆TM-OFP with failure to localize at the membrane was less efficient in reducing the accumulation of BaMV.

The results of yeast two-hybrid and co-immunoprecipitation assays revealed that the RdRp domain of BaMV replicase is the substrate of NbUbE3R1. We are missing the link of how and where NbUbE3R1 targets BaMV replicase. The availability of BaMV replicase for targeting could be in chloroplasts, where viral RNA replication occurs, or in the cytoplasm, where replicase is translated, or in the membrane-associated movement complex. Although NbUbE3R1 containing the transmembrane domain and the membrane-associated character was most efficient in reducing BaMV replication, a small portion of NbUbE3R1 transported to the chloroplasts could not be excluded.

The E3 ubiquitin ligases of the UPS were found involved in disease resistance as regulators of the plant defense responses (Azevedo *et al.*, 2002; Berrocal-Lobo *et al.*, 2010; Verchot, 2014). The replication enzyme p92 of *Tomato bushy stunt virus* (TBSV) was found regulated by the UPS (Barajas *et al.*, 2009; Li *et al.*, 2008). The ubiquitination of TBSV p33 could interrupt the interaction with the host protein Vps23p endosomal complexes required for transport (ESCRT), with failure in viral RNA replication (Barajas and Nagy, 2010). Also, the 66-kDa RdRp of TYMV was targeted by the UPS in infected plant cells and downregulated viral replication (Camborde *et al.*, 2010).

The UPS could also affect viral cell-to-cell movement. Increasing evidence has shown that viral MPs are regulated by proteasome turnover. The 69-kDa MP of TYMV could be polyubiquitinated and degraded by proteasome (Drugeon and Jupin, 2002; Verchot, 2014). The 17-kDa MP of *polerovirus*, TGB3 of PVX, and the 30-kDa MP and CP of *Tobacco mosaic virus* were regulated by the UPS degradation pathway (Jockusch and Wiegand, 2003; Ju *et al.*, 2008; Mas and Beachy, 1999; Pazhouhandeh *et al.*, 2006; Verchot, 2014). The CP of HIV type 1 was targeted by TRIM5 (the tripartite motif protein), which exhibits E3 ubiquitin ligase activity, and was removed by proteasome-dependent degradation (Li *et al.*, 2013). This type of protein turnover may represent an antiviral defense mechanism (Hwang *et al.*, 2011). These studies suggest that the UPS may target viral protein to interfere in viral invasion. In this study, we have identified an E3 ubiquitin ligase in *N. benthamiana* which has not been reported before could target the replicase of BaMV and resulted in downregulating BaMV replication.

## Supplementary materials

**Supplementary Figure S1.**Expression profile of *ACGT2-1* in BaMV-infected *N. benthamiana* verified by semi-quantitative RT-PCR. Mock- and BaMV-inoculated samples are indicated as M and I, respectively. The samples were harvested at 1, 3, 5 and 7 days post inoculation indicated above each lane. The expression of actin was an internal control.

**Supplementary Figure S2.**Structural features of NbUbE3R1. Illustration of NbUbE3R1 structure. I: low complexity region; II: transmembrane region; III: RING finger domain; and IV: coiled coil region.

**Supplementary Figure S3.**Phenotype of control and *NbUbE3R1*-knockdown plants. Phenotype of *Agrobacterium*-mediated knockdown plants at 14 days post infiltration. Phytoene desaturase gene-knockdown plants exhibiting a photobleach phenotype were the positive control. Morphology of *NbUbE3R1*-knockdown plants did not differ from that of *luciferase*-knockdown plants, the negative control.

## Acknowledgements

We thank the Bioimage core lab of the Graduate Institute of Biotechnology at National Chung Hsing University for use of the confocal laser scanning microscope.

